# Both ATP and Mg^2+^ are Required for High-Affinity Binding of Indolmycin to Human Mitochondrial Tryptophanyl-tRNA Synthetase

**DOI:** 10.64898/2026.03.23.713518

**Authors:** Charles W. Carter

**Author notes:** To whom correspondence should be addressed: Charles W. Carter, Jr Department of Biochemistry and Biophysics, University of North Carolina at Chapel Hill, Chapel Hill, NC 27599-7260 USA.

## Abstract

Eukaryotes have distinct nuclear genes for tryptophanyl-tRNA synthetase (TrpRS). Human mitochondrial (H_mt_) TrpRS (also WARS2) shares only 14% sequence identity with human cytoplasmic (H_c_)TrpRS, but 41% with *Bacillus stearothermophilus* (Bs)TrpRS. Tryptophan binding to BsTrpRS is largely promoted by hydrophobic interactions and recognition of the indole nitrogen by side chains of Met^129^ and Asp^132^. The non-reactive analog indolmycin can recruit unique polar interactions to form an active-site metal coordination that lies off the normal mechanistic path, enhancing affinity to BsTrpRS and other prokaryotic TrpRS enzymes by 1500-fold over its tryptophan substrate. By contrast, human WARS2, complements nonpolar interactions for tryptophan binding with additional electrostatic and hydrogen bonding interactions that are inconsistent with indolmycin binding. We report here a 1.82 Ȧ crystal structure of an H_mt_TrpRS• indolmycin•Mn^2+^•ATP complex, showing that mitochondrial and bacterial enzymes use similar determinants to bind both ATP and indolmycin. ATP forms tight electrostatic interactions between the catalytic metal ion and a non-bridging oxygen atom from each phosphate group. Hydrogen bonds between the oxazolinone group and active-site residues create an off-path ground-state configuration. This arrangement closely mimics that in the corresponding BsTrpRS complex but varies greatly from ATP binding to H_c_TrpRS, Moreover, isothermal titration calorimetry demonstrates that, as for BsTrpRS, Mg^2+^•ATP, but not ATP alone, enhances indolmycin binding affinity ∼100-fold with a supplemental Δ(ΔG) of ∼ -3 kcal/mol. Structural, thermodynamic, and kinetic similarities confirm our previous conclusion that a reinforced ground-state Mg^2+^ ion configuration contributes to the high indolmycin affinity in the bacterial system.

## INTRODUCTION

Increasing drug resistance places increasing demands on the development of antibiotics active against novel targets. Mitochondrial dysfunction is often a detrimental consequence of prolonged antibacterial and antiviral therapeutics due, in part, to unintended drug interactions with host mitochondrial homologs of the pathogenic target proteins (1-4). Further, mutations to mitochondrial aminoacyl-tRNA synthetases have recently been linked to various chronic neurologic diseases

Mitochondria have retained many conserved proteins, pathways and processes from their prokaryotic ancestors. Dysregulation or inactivation of mitochondrial enzymes, most of which are nuclear-encoded, has great potential for debilitating ATP production. Successful pharmaceutical lead candidates therefore should be assessed for off-target effects including the potential for deleterious interactions with mitochondrial proteins, which are often closely related to bacterial homologs.

One under-utilized target for antibacterial action is tryptophanyl-tRNA synthetase. We previously demonstrated that mechanistic differences allow indolmycin, a natural TrpRS inhibitor, to selectively bind *Bacillus stearothermophilus* (Bs) TrpRS 10^6^-fold over human cytosolic (H_c_) TrpRS (5). Specifically, BsTrpRS uses a combination of interactions between Met^129^ and an acidic residue, Asp^132^, to recognize the indole nitrogen and bind indolmycin (and substrate tryptophan) to a largely hydrophobic pocket. A catalytic Mg^2+^ ion electrostatically coupled to each phosphate group of ATP assumes a high energy configuration that we have proposed activates it for participation in catalysis (6-8).

The methylamino-substituted oxazolinone ring (OXA) of indolmycin prevents Mg^2+^ activation by forming a more stable, hexavalent Mg^2+^ coordination, resulting in stronger Mg^2+^•ATP interactions, and weaker TrpRS•ATP interactions (5). Sequence variations, specifically alanine substitution for the second lysine of the conserved KMSKS motif and shifting of catalytically-important residues to the N-terminal extension, alter the ATP configuration and Mg^2+^•ATP coordination in H_c_TrpRS. These differences and a greater number of polar binding determinants for tryptophan recognition prevent indolmycin from binding tightly (9-11).

Although the human mitochondrial (H_mt_) TrpRS enzyme was identified and kinetically characterized over 15 years ago, structural information about the enzyme remains lacking (12). Thus, little is known about how the enzyme recognizes its substrates. Based solely on primary sequence comparison, H_mt_TrpRS has retained the aspartic acid residue of the specificity helix shown to recognize tryptophan in BsTrpRS while showing bacteriospecific variations at other positions involved in tryptophan binding (Fig. 1). Additionally, H_mt_TrpRS has the canonical KMSKS sequence important for ATP and pyrophosphate binding and lacks an N-terminal extension acquired by H_c_TrpRS (Fig. 1). From these differences, we expect H_mt_TrpRS to utilize similar determinants as BsTrpRS for tryptophan and ATP binding and, similarly, to be sensitive to inhibition by indolmycin.

**Figure 1:**
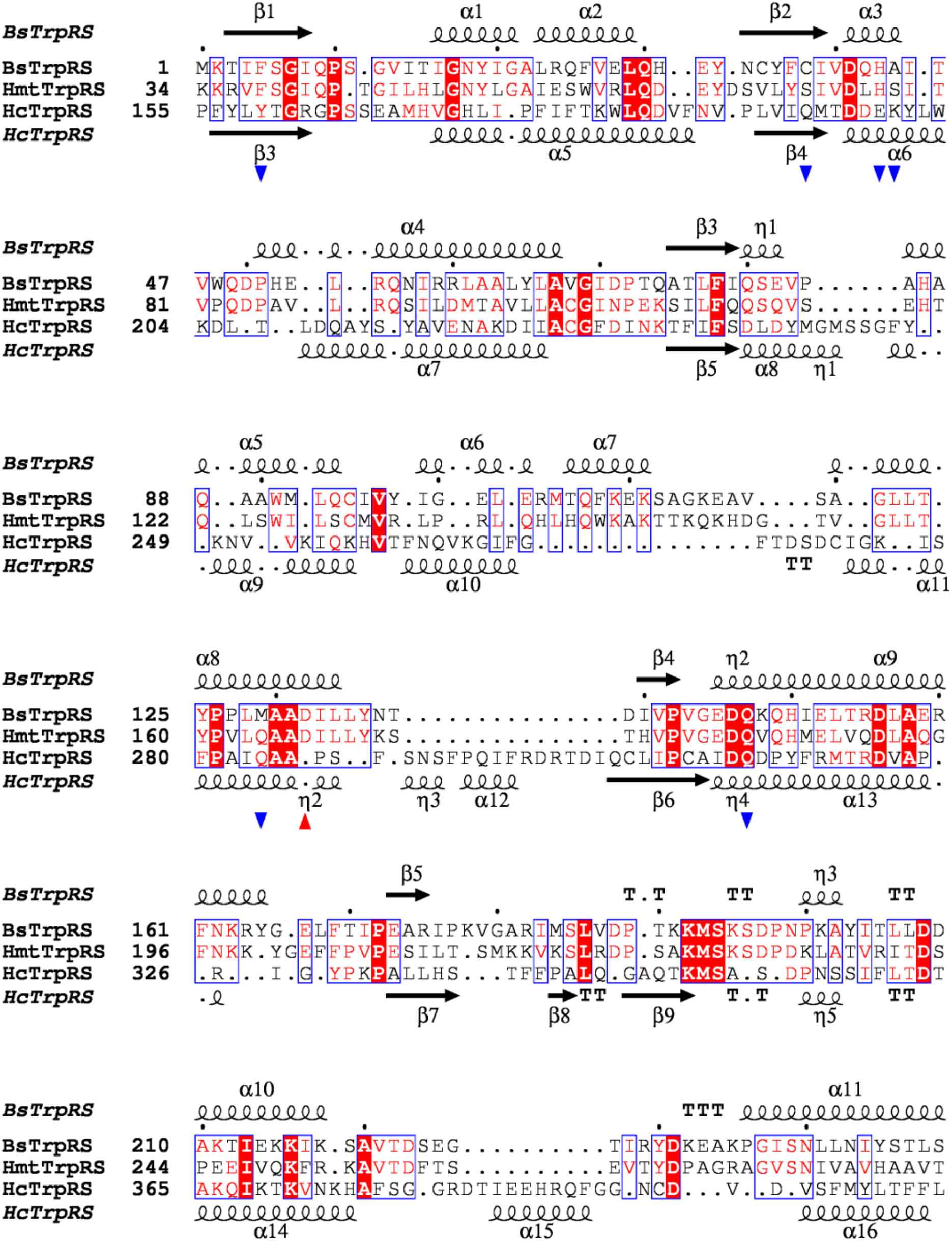
Structure-based sequence alignment of BsTrpRS, H_mt_TrpRS and H_c_TrpRS. Following superposition of relevant BsTrpRS, H_mt_TrpRS and H_c_TrpRS structures USCF Chimera was used to generate a structure-based sequence alignment of the three enzymes. This alignment facilitates comparison of residues used for substrate binding. The red and blue triangles point to residues that interact with tryptophan when tryptophan alone is bound to BsTrpRS and H_c_TrpRS, respectively. The final image was generated using ESPript 3.0.

Here, we provide structural, kinetic and thermodynamic evidence that indolmycin also is a tight-binding inhibitor of H_mt_TrpRS. Additionally, we use the crystal structure of H_mt_TrpRS determined as part of this work to describe structural details that support a shared mechanism for indolmycin inhibition between H_mt_TrpRS and BsTrpRS as well as utilization of conserved elements in the recognition/binding of ATP and indolmycin that imply a common mechanism for tryptophan activation. Notably, isothermal titration calorimetry reveals that indolmycin binding affinity is strengthened by simultaneous binding of Mg^2+^•ATP, but not ATP.

## EXPERIMENTAL PROCEDURES

### Construction of pet28-His-H_mt_TrpRS

A gene containing residues 19-360 of H_mt_TrpRS was purchased from GenScript. A tobacco etch virus (TEV) cleavage site, ENLYFQG, was added to the N-terminus of H_mt_TrpRS by PCR. The resultant PCR product was digested with restriction enzymes NdeI and HindIII and ligated into pet28b. This yielded an expression vector for His-tev-H_mt_TrpRS.

### Expression and Purification of His-H_mt_TrpRS

Production of 6xHis-H_mt_TrpRS was performed in E. coli BL21(DE3) containing the vectors pet28-His-H_mt_TrpRS and pGro7 (13). Overnight cultures were grown in ZYM-505 (14) with 100 μg/ml kanamycin and 25 μg/ml chloramphenicol. Arabinose (0.05%) was used to induce the GroEL/ES chaperones. In a 4L flask, 1L of ZYM-5052 auto-inducing media with 0.05% arabinose was inoculated with the overnight culture to a starting OD_600_ of 0.05. Cells were grown at 37°C until the culture reached an OD_600_ of 1, at which time the temperature was shifted to 18°C and allowed to grow for ∼16 hours. The cells were pelleted at 4500 rpm for 30 minutes, resuspended in lysis buffer and frozen at -20°C. Upon thawing, cells were sonicated and centrifuged (16000 rpm, 4°C, and 1 hour). His-H_mt_TrpRS was captured from the lysate on Ni-NTA resin and eluted with 0.3 M imidazole. Purified protein was cleaved overnight with tobacco etch virus (TEV) while dialyzing out the imidazole. The cleaved protein mixture was passed back over a Ni-NTA column to capture both uncleaved protein and his-TEV protease. Size exclusion chromatography on a Superdex 200 column was used as a final purification step for H_mt_TrpRS.

### Active Site Titration

Active sites were titrated by following the loss of [γ-^32^P] Type equation here. ATP to determine the fraction of molecules competent for catalysis as described by (15, 16). Plates were developed using a Typhoon Imager and analyzed with ImageJ (17) and JMP (18).

### Crystallization, Data Collection, Structure Determination

Crystals of H_mt_TrpRS in complex with ATP, indolmycin and Mn^2+^ were grown by vapor diffusion against 25% glycerol, 20% Peg5000 and 0.1 M sodium acetate. As the crystal growth solution was cryoprotective, crystals did not require further handling before plunging into liquid nitrogen. Data were collected at Southeast Regional Collaborative Access Team (SER-CAT) 22-ID beamline at the Advanced Photon Source, Argonne National Laboratory. Supporting institutions may be found at www.ser-cat.org/members.html. The structure was determined using molecular replacement with the BsTrpRS structure (5DK4) as a starting model’. The initial model was improved using cycles of Coot (19) rebuilding and refinement using Phenix (20).

### Michaelis-Menten Kinetics

Incorporation of [^32^P]PPi into ATP was tracked by thin-layer chromatography. Reactions contained 5000 cpm/μl [^32^P]PPi, 0.1 M Tris pH 8.0, 70 mM 2-mercaptoethanol, 5 mM MgCl_2_, 10 mM potassium fluoride, 2 mM pyrophosphate, 2 mM ATP and tryptophan ranging from 1-500 μM. When ATP was varied from 60-4000 μM tryptophan was used at a final concentration of 2 mM. Reactions were initiated with enzyme.

### Indolmycin Inhibition Assays

Inhibition assays were performed as described above; Michaelis-Menten experiments for tryptophan were performed in the presence of stoichiometric amounts of indolmycin to enzyme. Indolmycin to enzyme ratios of 1:5, 1:1 and 5:1 were used and results fitted to a competitive inhibition model (I) using JMP. This setup allowed simultaneous determination of K_M_ for tryptophan and K_i_ for indolmycin.

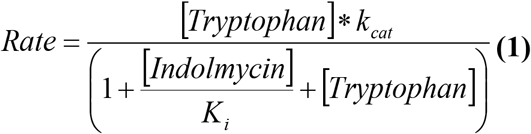

### Isothermal Titration Calorimetry

Experiments were carried out at 37°C. The sample cell was filled with H_mt_TrpRS in 50 mM Na_2_HPO4, 150 mM NaCl, 10 mM 2-mercaptoethanol, pH 7.4 with and without 10 mM MgCl_2_ and 5 mM ATP. Using the MicroCal Auto-iTC200 system, ligands were added by a series of 20-2 μl injections with 180 seconds between each injection and a reference power of 8 μcal/sec. Twenty-seven 1.5 μl injections were used to titrate H_mt_TrpRS pre-bound to Mg^2+^•ATP with indolmycin. Data were analyzed using a one-site model in Origin *(OriginLab, Northampton, MA)*.

## RESULTS

Justesen *et al*. (12) described residues 1-19 as the mitochondrial targeting signal, while MitoProt (http://ihg.gsf.de/ihg/mitoprot.html) predicted that the localization signal continues through residue 33. Michaelis-Menten kinetics for full-length, Δ 1-18, and Δ 1-33 variants of H_mt_TrpRS showed comparable k_cat_/K_M_ values (Table 1). As the Δ 1-18 variant consistently yielded higher protein concentrations and was efficiently cleaved by TEV protease, all kinetic, thermodynamic and structural studies were carried out with this variant.

**Table 1:** Data collection and refinement statistics for H_mt_TrpRS inhibited complex. The H_mt_TrpRS•Indolmycin•Mn^2+^•ATP complex was determined to 1.82 Å resolution by molecular replacement using the BsTrpRS inhibited complex (PDB 5DK4) as a model. The final structure was refined to an R_Work_/R_Free_ of 17.0/20.7%.

| H<sub>mt</sub>TrpRS | k<sub>cat</sub> (s<sup>-1</sup>) | K<sub>M</sub> (M) | k<sub>cat</sub>/K<sub>M</sub> (s<sup>-1</sup> M<sup>-1</sup>) | (k<sub>cat</sub>/K<sub>M</sub>)<sub>relative</sub> |
| --- | --- | --- | --- | --- |
| 1-360 | 3.17 ± 0.02 | 2.35E-04 ± 9.07E-06 | 1.35E04 ± 4.57E02 | 1.00 ± 0.05 |
| Δ 1-18 | 4.61 ± 0.03 | 3.59E-04 ± 2.23E-05 | 1.29E04 ± 7.34E02 | 0.96 ± 0.06 |
| Δ 1-33 | 3.31 ± 0.21 | 2.33E-04 ± 1.00E-06 | 1.42E04 ± 9.01E02 | 1.05 ± 0.08 |

### Indolmycin binds to H_mt_TrpRS as a ternary complex with ATP

Unlike the closed-state crystal structures of BsTrpRS, all of which have a monomer in the asymmetric unit, H_mt_TrpRS crystallized in space group C121 with a dimer in the crystallographic asymmetric unit and crystals diffracted to 1.82 Å (Fig. 2a and Table 2). Each monomer contains ATP, Mn^2+^ and indolmycin in the active site at essentially full occupancy. Mn^2+^ was used instead of Mg^2+^ because we hoped to use the Mn^2+^ anomalous signal to derive experimental phase information. Although the signal was too weak for phasing, the anomalous signal did allow us to definitively identify the metal binding site. There was no electron density in which to place N-terminal leader sequences, residues 19-33 and 19-28 in monomers A (grey) and B (pink), respectively.

**Table 2:** Catalytic efficiency of H_mt_TrpRS is independent of residues 1-33. Steady-state kinetics experiments tracking the incorporation of [^32^P]PPi into ATP were performed at saturating tryptophan and variable ATP concentrations with three variants of the H_mt_TrpRS enzyme. The data, fitted by non-linear regression modeling to simultaneously determine K_M_ ATP and k_cat_, show that the three variants have statistically indistinguishable catalytic efficiencies. ^a^Highest resolution shell is shown in parentheses.

| **Data Collection** |  |
| --- | --- |
| Space group | C 1 2 1 |
| Cell Constants | 58.289 Å 78.087 Å 153.996 Å |
| a, b, c, $\alpha$, $\beta$, $\gamma$ | 90.00° 96.55° 90.00° |
| Resolution (Å) | 29.06 – 1.82 |
| Completeness (%)<sup>a</sup> | 98.5 (95.8) |
| Rmerge (%)<sup>a</sup> | 7.7 (59.4) |
| Rmeas (%)<sup>a</sup> | 9.8 (76.1) |
| Rpim (%)<sup>a</sup> | 6.0 (47.0) |
| Mean I/σI<sup>a</sup> | 8.1 (1.5) |
| Number of Reflections | 139667 |
| Multiplicity | 2.3 |
| **Refinement** |  |
| R<sub>work</sub>/R<sub>free</sub> (%) | 17.2/20.7 |
| Fo, Fc correlation | 0.96 |
| RMS (bonds) | 0.004 |
| RMS (angles) | 0.92 |
| Ramachandran favored (%) | 97.0 |
| Ramachandran outliers (%) | 0.0 |
| Average B, all atoms (Å<sup>2</sup>) | 28.8 |
| Clash score | 5.76 |
| PDB Entry ID | 5EKD |

<sup>a</sup>Highest resolution shell is shown in parentheses.

**Figure 2:**
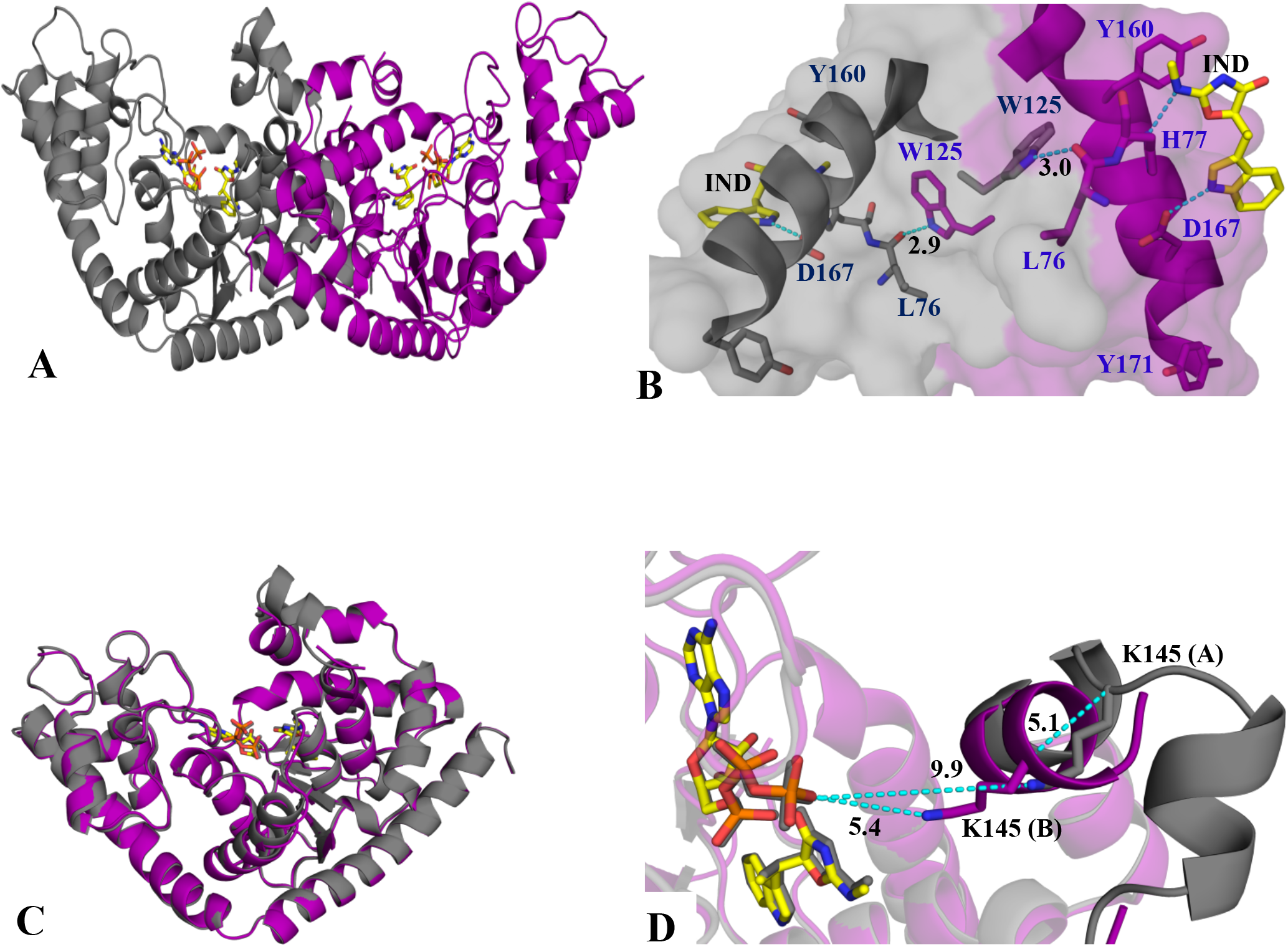
Dimeric H_mt_TrpRS binds one molecule of ATP and indolmycin per active site. (A)Two monomers, A (grey) and B (purple), form the functional dimer. (B) A prominent feature of the dimer interface is the inter-monomeric burial of Trp^125^, whose side chain is stabilized by a hydrogen bond to the carbonyl oxygen of Leu^76^. This feature was also observed in several BsTrpRS structures. (C) Structurally, the two monomers are quite similar but show variation in the position of residues 140-150. (D) The importance of this difference to ligand binding is unknown. However, Lys^145^ is analogous to BsTrpRS Lys^111^, which shows ligand-dependent structural variation.

Using areaImol (21, 22), monomer A has a solvent-accessible area of 13957.4 and 15918.3 Å^2^ in the presence and absence of monomer B, respectively. The solvent-accessible area for monomer B alone is 15642.4 Å^2^ and 13695.5 Å^2^ in the dimer. The calculated buried surface area due to dimer formation is 3907.8 Å^2^. Similar calculations using pymol yielded a buried surface area of 3815.5 Å^2^. As is the case for the bacterial enzyme, a prominent feature of the dimer interface is the burial of a tryptophan side-chain (Trp^125^) across the dimer interface into a space bounded by the specificity helix of the other monomer (Fig. 2b). A hydrogen bond between the carbonyl oxygen of Leu^76^ and the NE1 of Trp^125^ helps stabilize the dimer interface. As measured from CH2, Trp^125^ is 4.0 Å away from OG1 of Thr^159^ and 4.1 Å from the CA of Tyr^160^. Both Thr^159^ and Tyr^160^ are part of the specificity helix and Tyr^160^ stacks against the oxazolinone ring of indolmycin.

Superposition of the two monomers reveals structural differences in the region of loop 144-150 (Fig. 2c,d) that are presumably responsible, in part, for the inclusion of a dimer in the asymmetric unit. This loop is also difficult to position in BsTrpRS structures (23) and is responsible for much of the conformational differences within the catalytic domain along the structural reaction profile (24) (25). As noted below, this loop thus probably has an important role in transition state stabilization either in amino acid activation, acyl-transfer, or both. It is disordered in H_mt_TrpRS monomer B, as evidenced by a lack of electron density, but well-ordered in monomer A. Residues His^140^ to Ala^144^ of monomer A are helical and angled away from the active site compared to the helix formed by His^140^-Thr^146^ in monomer B. Residues from Thr^147^ to Asp^152^ form a helix in monomer A but cannot be placed in monomer B. This region contains a catalytically-relevant lysine residue, Lys^145^, which is homologous to Lys^111^ in BsTrpRS.

Loop 148-152 corresponds to residues 112-117 in BsTrpRS, which also was fitted with difficulty for some structures, depending on which ligands occupied the active site. The relative positions of α-carbon atoms of Lys^145^ in the two monomers differ by 5.1 Å following superposition of monomer B onto monomer A (Fig. 2d). The NZ atom of Lys^145^ is 9.9 Å away from the nearest gamma phosphate oxygen atom in monomer A but only 5.4 Å away in monomer B. While neither residue is in position to form direct interactions with ATP as is observed in the BsTrpRS pre-transition state and inhibited complexes (5, 23) Lys^145^ in monomer B does structurally align with BsTrpRS Lys^111^, and it interacts with the gamma phosphate of ATP via a water molecule.

Kinetic analysis of mutants in this region in both TrpRS (7, 26) and TyrRS (26) show that residues from this loop contribute to ATP binding only in the transition state and imply that movement of the loop occurs during catalysis. In BsTrpRS Lys^111^ competes with the catalytic Mg^2+^ ion for the same negatively charged gamma phosphate oxygen atom in the pre-transition state (2.9 Å) and inhibited state (3.4 Å) complexes. Mutation of Lys^111^ to alanine affects k_cat_ but not K_M_ for ATP (7).

The H_mt_TrpRS anticodon-binding domain is rotated ∼5 degrees closer to the Rossmann fold domain than it is in the PreTS and inhibited BsTrpRS structures. Without knowing structures of tRNA complexes for either protein, it is possible only to speculate that this difference relates to differences between the respective tRNA structures, and note that the mitochondrial tRNAs and tRNATrp in particular have shortened acceptor-TΨC stems that would bring the anticodon closer to the acceptor stem (27, 28).

### Indolmycin Interactions

The indole moiety of indolmycin is recognized by the combination of Asp^167^ and Gln^170^ in the specificity helix, as is observed for homologous interactions with Asp^132^ and Met^129^ in BsTrpRS complexes (5) (Fig 2b). Specifically, the sidechain of Asp^167^ accepts a hydrogen bond (3.0 Å) from the pyrrole ring nitrogen and the amide π electron system of Gln^170^ interacts with the five-membered ring of indole. Phe^38^ forms π-π stacking interactions while Ile^41^, Val^74^, Ile^168^, Val^176^ and Val^178^—all hydrophobic—make van der Waals contacts with indolmycin. Sidechain atoms of His^77^ and Gln^182^ donate hydrogen bonds to the methylamino nitrogen (3.0 Å) and carbonyl oxygen (3.1 Å) atoms of the oxazolinone moiety, respectively, restraining its orientation relative to Mg^2+^•ATP. The carbonyl oxygen accepts an additional hydrogen bond from a metal-coordinating water molecule, as does the oxazolinone nitrogen (Fig. 3a, b).

**Figure 3:**
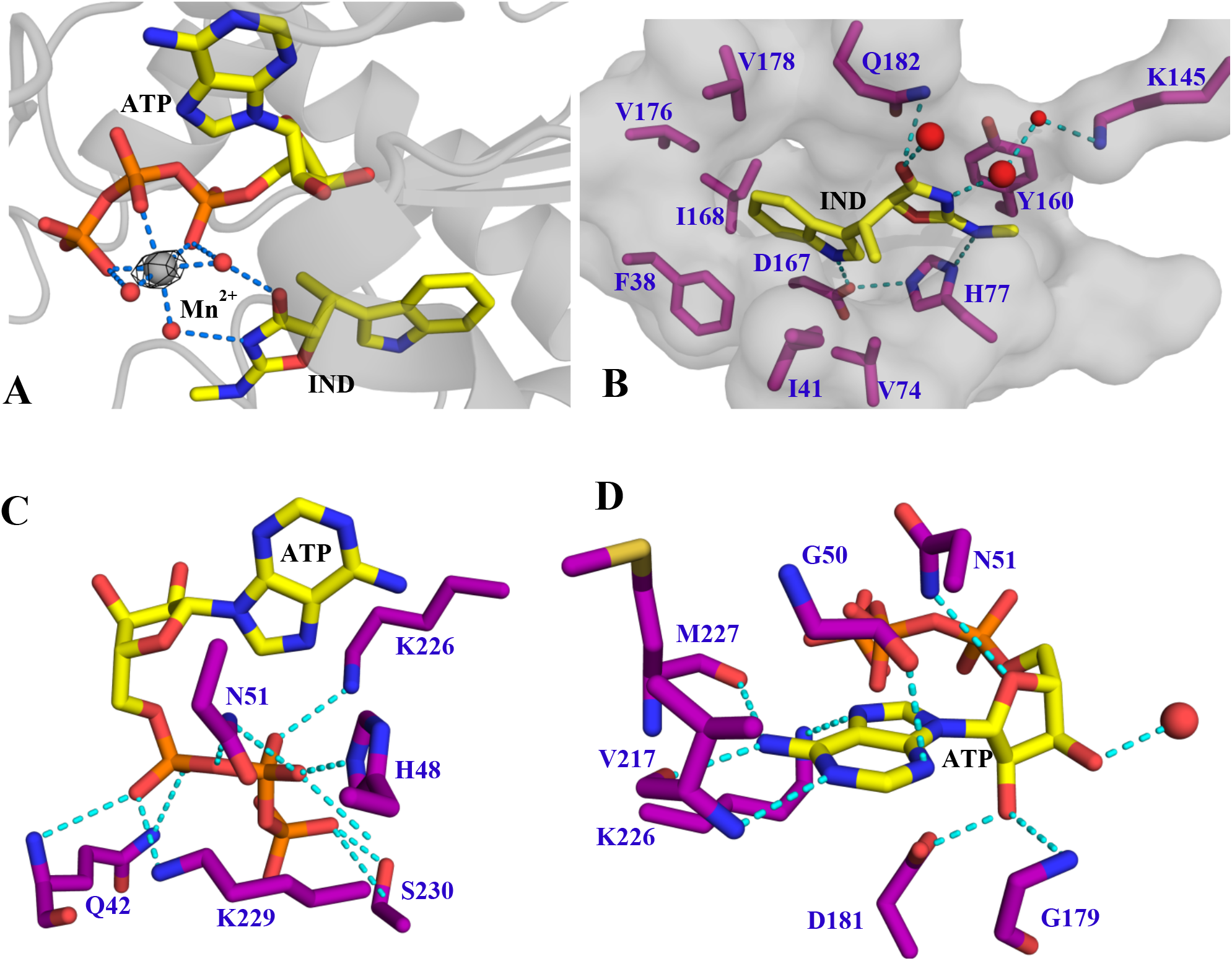
Molecular determinants of indolmycin and ATP binding are conserved in H_mt_TrpRS and BsTrpRS. (A) Like BsTrpRS, the inhibited H_mt_TrpRS complex involves a hexacoordinated-metal ion. Bijvoet difference fourier map due to manganese is depicted in blue mesh contoured to 5σ. (B) Binding of indolmycin involves hydrophobic, π-π, and hydrogen bonding interactions. The interactions with His^77^, Asp^167^, Gln^182^ and two metal-coordinated water molecules mirror those observed in the inhibited BsTrpRS complex. (C) ATP binds in an extended configuration with Gln^42^ and residues of the HIGH and KMSKS motifs stabilizing the triphosphate moiety. (D) The adenine and ribose moieties of ATP are recognized by the three signature sequences for class I aaRS enzymes; HIGH, GxDQ and KMSKS.

### ATP Interactions

H_mt_TrpRS binds ATP in an extended conformation and utilizes conserved residues for substrate recognition and binding. As was observed for the BsTrpRS•ATP•indolmycin complex, the negative charges of the triphosphate oxygen atoms form strong electrostatic interactions with a metal ion. A water molecule coordinated by the ATP 3’OH group, the hydroxyl group of Ser^39^ and the backbone carbonyl of Gly^40^ represents an additional detailed homology with the BsTrpRS PreTS (23), Product (29), and inhibited (5) complexes.

The Mn^2+^ is hexavalently coordinated by an oxygen atom from each phosphate group and three water molecules (Fig. 3a). The three phosphate groups are additionally stabilized by direct and water-mediated protein interactions. Gln^42^ (2.8-3.0 Å), Asn^51^ (3.1 Å) and Lys^229^ (2.8 Å) form contacts with P_α_, while Lys^226^ (2.9 Å), Lys^229^ (2.9 Å) and His^48^ (2.9 Å) donate hydrogen bonds to P_β_ oxygen atoms (Fig. 3c). The backbone amide and sidechain hydroxyl groups of Ser^230^ contribute to the stabilization of Pγ via two hydrogen bonds (3.2 and 2.6 Å). Thr^44^, Lys^145^ (B), Ser^228^, and Lys^229^ form additional water-mediated interactions with gamma-phosphate oxygen atoms.

The adenine 6-amino group donates a hydrogen bond to the backbone carbonyl oxygen atoms of Val^217^ (2.9 Å) and Met^227^ (2.9 Å) (Fig. 3d). The nitrogen atoms of the adenine ring accept hydrogen bonds from the amide groups of Val^217^ (3.1 Å) and Gly^50^ (3.1 Å) as well as from the sidechain NZ atom of Lys^226^ (3.0 Å). The ribose moiety interacts with the conserved HIGH (48-51) and GxDQ (179-182) motifs. Specifically, the 2’OH group accepts a hydrogen bond from Gly^179^ (3.0 Å) and donates a hydrogen bond to Asp^181^ (2.7 Å) while Asn^51^ donates a hydrogen bond to the ringed oxygen atom. The 3’OH group forms a close interaction (2.6 Å) with a water molecule (Fig. 3d). These interactions mimic closely those observed in BsTrpRS pre-transition state complexes (6, 23, 30).

### Indolmycin is a tight-binding inhibitor of H_mt_TrpRS

Table 3 summarizes our determination of the indolmycin inhibition constant by steady-state kinetics. The human mitochondrial (H_mt_)TrpRS enzyme is about 130-fold less proficient than BsTrpRS, owing to roughly similar differences in both k_cat_ (7-fold smaller) and K_M_ (19-fold greater). Despite the extensive structural similarity between indolmycin and substrate tryptophan both BsTrpRS and H_mt_TrpRS selectively bind indolmycin tighter than the natural tryptophan substrate by 2 to 3 orders of magnitude. In contrast, cytosolic isoforms of eukaryotic TrpRS bind tryptophan ∼1000x tighter than indolmycin.

**Table 3:**
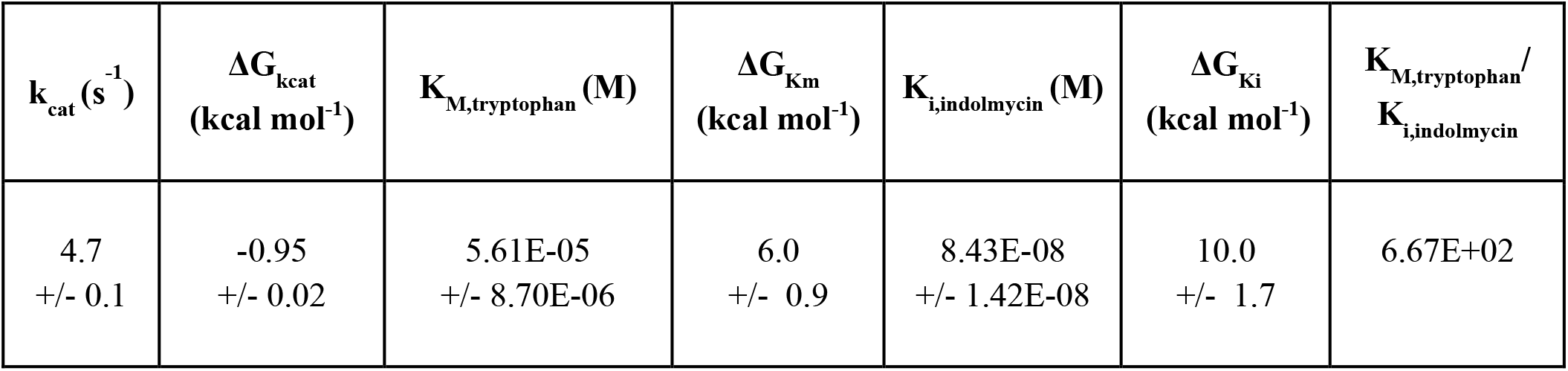
Indolmycin is a competitive inhibitor of H_mt_TrpRS. Steady-state kinetics experiments performed with saturating amounts of ATP at various tryptophan and indolmycin concentrations show indolmycin inhibits H_mt_TrpRS with nanomolar affinity.

Thus, although both substrate and inhibitor share the indole ring and have similar hydrogen bonding groups associated with all three α-carbon substituents, indolmycin exploits apparently minor differences elsewhere to enhance binding to bacterial TrpRS, yet reduce binding to the cytosolic TrpRS, leading to a selectivity between the cytosolic and bacterial forms of close to 10^6^-fold. From our studies we determined that H_mt_TrpRS is 42-times less likely to bind indolmycin than is BsTrpRS and ∼10^4^-times more likely than eukaryotic cytosolic TrpRS enzymes. The ∼40-fold reduced sensitivity of mitochondrial TrpRS relative to BsTrpRS defines a quantitative window within which it may be possible to design useful bacterio-specific anti-infective agents.

### Mg^2+^•ATP is required for high-affinity binding of indolmycin to H_mt_TrpRS

Our previous study on indolmycin inhibition of BsTrpRS revealed that while indolmycin enhanced the thermal stability of BsTrpRS by 13.5°C, addition of Mg^2+^•ATP to form the fully-liganded inhibited complex resulted in a further thermal stabilization of 5°C (5). The metal ion was critical for this effect, as ATP alone did not alter the thermal stability over that observed for indolmycin alone. We hypothesized that high-affinity indolmycin inhibition of bacterial TrpRS resulted, in part, from locking the Mg^2+^ ion into a ground state configuration that precludes catalysis. The overwhelming similarities in ligand (ATP, metal ion, indolmycin) arrangement and coordination within the active sites of BsTrpRS and H_mt_TrpRS lead to the following predictions: 1) Mg^2+^ alone should not alter indolmycin binding—since the metal ion only forms direct contacts with ATP and water molecules, it should not bind in the absence of ATP; 2) ATP alone may alter the affinity of H_mt_TrpRS for indolmycin—in addition to interacting through Mg^2+^, the two ligands are linked via the conserved GxDQ motif with Gly^179^, Asp^181^ and Gln^182^ forming hydrogen bonds with the 2’OH group of ATP and carbonyl oxygen of indolmycin; and 3) Mg^2+^•ATP should enhance the affinity of H_mt_TrpRS for indolmycin—the presence of both ATP and indolmycin allow Mg^2+^ to fall into a stable position inconsistent with catalysis, thereby contributing to high-affinity inhibition by indolmycin.

We employed isothermal titration calorimetry (ITC) to test predictions 2 and 3 using indolmycin as the titrant for ligand-free, ATP-bound, and Mg^2+^•ATP-bound H_mt_TrpRS (Fig. 4a, b, c). In all cases we observed 1:1 binding of indolmycin to H_mt_TrpRS. As we used the concentration of active sites, determined by active site titration prior to ITC, in our analysis this means indolmycin bound to each active site of the functional dimer. The indolmycin dissociation constant (K_d_) and Gibbs free energy of binding (ΔG) are 3.9 ± 0.3 μM and -7.7 ± 0.1 kcal/mol, respectively (Fig. 4a, 5b). In the absence of Mg^2+^, pre-binding with ATP led to a three-fold increase in indolmycin affinity (Fig. 4b). The H_mt_TrpRS•ATP•IND complex (ΔG = -8.4 ± 0.1 kcal/mol) is more stable than the H_mt_TrpRS•IND complex by -0.7 kcal/mol (Fig. 5b). Addition of both Mg^2+^ and ATP to form the fully-liganded inhibited complex enhances the affinity by two orders of magnitude to 39.6 ± 8.4 nM, a value that is similar to our kinetically-derived K_i_ of 84.5 ± 14.3 nM (Fig. 4c and Table 3). This complex (ΔG = -10.6 ± 0.3 kcal/mol) is more stable than either IND only or IND+ATP H_mt_TrpRS complexes by -2.9 and -2.2 kcal/mol, respectively (Fig. 5b).

**Figure 4:**
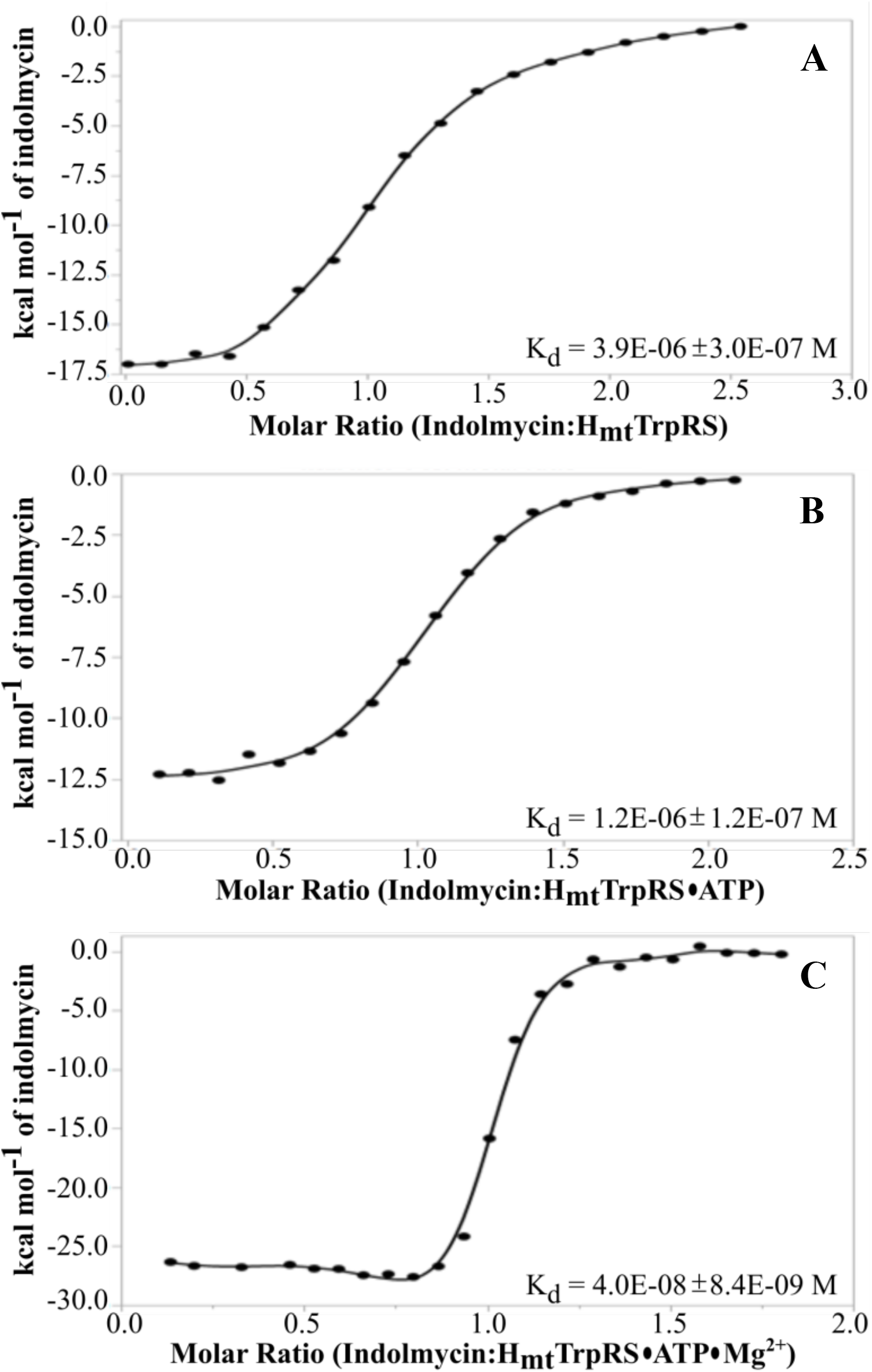
Mg^2+^ ATP is required for high-affinity binding of indolmycin to H_mt_ TrpRS. Isothermal titration calorimetry experiments revealed a 1:1 binding stoichiometry of indolmycin:H_mt_TrpRS active site. H_mt_TrpRS binds indolmycin with μM affinity in the absence (A) and presence (B) of ATP. Pre-binding with Mg^2+^•ATP (C) enhances indolmycin binding ∼100-fold.

**Figure 5:**
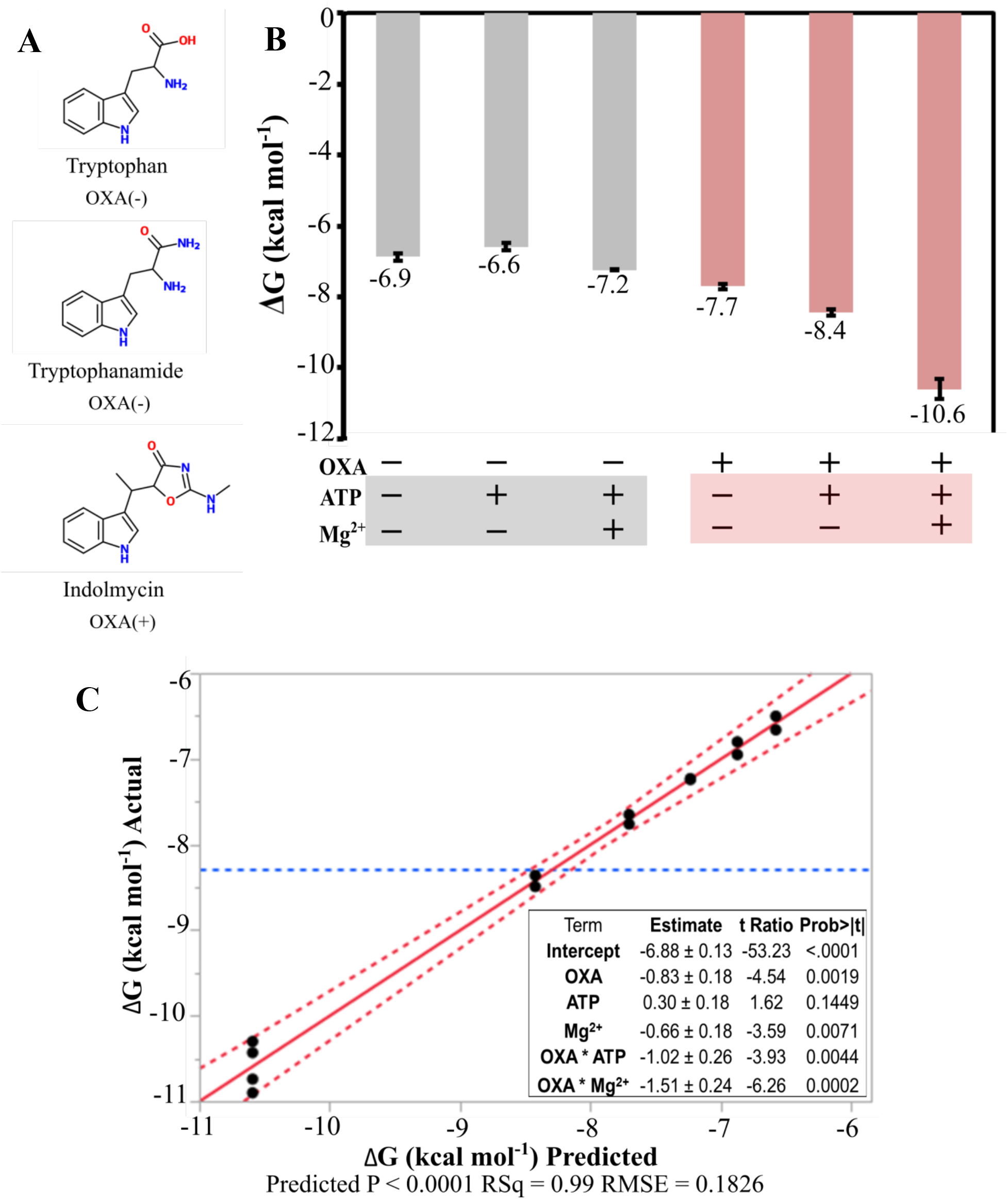
The potentiation effect of Mg^2+^•ATP is mediated via the methylamino-substituted oxazolinone (OXA) moiety of indolmycin. (A) Tryptophanamide is a conservative substitute for the tryptophan substrate in which the sole difference is replacement of a carboxylate oxygen atom by an amine group. Both have an indole moiety and lack the oxazolinone ring present in indolmycin. (B) The relative stability of six H_mt_TrpRS complexes were assessed using thermodynamic measurements obtained from replicate ITC experiments, in which ligand-free, ATP-, or Mg^2+^•ATP-bound H_mt_TrpRS were titrated with either tryptophanamide (OXA-) or indolmycin (OXA+). With a difference in Gibbs free energy of 4.0 kcal/mol the LTN•ATP and IND•Mg^2+^•ATP complexes are the least and most stable, respectively. (C) The ΔG values can be predicted using a model that contains five parameters which include the interaction energies of OXA with Mg^2+^ and ATP. The equation fitted is:

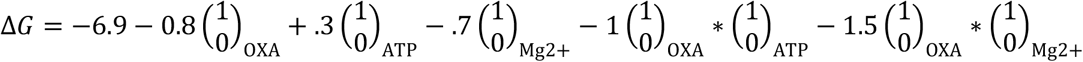

where 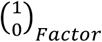 is an element of the design matrix and is either 1 or 0, according to whether the factor is present or not in that particular row, corresponding to a particular experiment.

### The methylamino-substituted oxazolinone moiety (OXA) is critical for the interaction between indolmycin and Mg^2+^•ATP

The heterocyclic indole moiety is common to both tryptophan and indolmycin with the major difference between these two compounds being the restriction of the Cα constituents via the oxazolinone ring (OXA) in indolmycin (Fig. 5a). We attribute differences in the thermodynamics of binding between tryptophan (OXA−) and indolmycin (OXA+) to the OXA moiety. Given our result that Mg^2+^•ATP stabilizes by ∼-3 kcal/mol binding of indolmycin to form the fully-ligated H_mt_TrpRS•Mg^2+^•ATP•IND complex, we wanted to assess the individual contributions of ATP, Mg^2+^ and the OXA moiety to high-affinity indolmycin binding and uncover any Gibbs coupling free energies (ΔΔG_int_) between these three components. Because we expected Mg^2+^ to bind H_mt_TrpRS only when ATP is also bound, we only examined the ATP-dependent effect of Mg^2+^ on ligand binding.

Using ITC, we were able to quantify the effects of ATP, Mg^2+^ and Mg^2+^•ATP on the Gibbs free energy of tryptophanamide (OXA−) binding to H_mt_TrpRS (Fig. 5b). Tryptophanamide (LTN), shown to bind TrpRS in a manner comparable to tryptophan, is a non-reactive tryptophan analog that allows us to probe the energetics of indole binding separate from catalysis. Pre-binding ATP weakened tryptophanamide binding to H_mt_TrpRS by 0.3 kcal/mol, while the addition of both Mg^2+^ and ATP led to formation of a complex more stable by -0.3 kcal/mol (Fig. 5b). The Mg^2+^ ion contributes -0.6 kcal/mol to the ΔG of tryptophanamide binding when ATP is also bound, thereby compensating for the destabilizing effect of ATP alone.

Figure 5b highlights how the energetic contributions of ATP, Mg^2+^, and Mg^2+^•ATP depend on whether the indole-containing ligand has the OXA moiety (indolmycin) or not (tryptophanamide). This finding implies a coupling interaction between the OXA moiety of indolmycin with both ATP and Mg^2+^ in the fully-ligated H_mt_TrpRS•Mg^2+^•ATP•IND complex.

Indeed, linear regression modeling of the data using the five independent parameters OXA, ATP, Mg^2+^, the OXA*ATP interaction, and the ATP-dependent OXA*Mg^2+^ interaction predicts the binding free energy, ΔG, of all complexes with an R squared of 0.99 (Fig. 5c). Notably, ATP alone does not contribute significantly to the binding except via its interactions. Both the coefficients (>1 Kcal/mole) of the model and their Student t-test P-values (<0.005) underscore the statistical significance of the synergistic effects of OXA*ATP and OXA*Mg^++^.

The most stable complex is that which has OXA (IND), ATP, and Mg^2+^ (ΔG = -10.6 kcal/mol) (Fig. 5b). In the absence of ligand coupling, the effects of adding OXA (ΔΔG_2_ = - 0.8 kcal/mol) and Mg^2+^•ATP (ΔΔG_5_ = ΔΔG_1_ + ΔΔG_8_ = -0.4 kcal/mol) should be additive (Fig. 6 blue, purple, amber). In this case, we would expect the H_mt_TrpRS•Mg^2+^•ATP•IND complex to have ΔG = ΔG_LTN_ + ΔΔG_2_ + ΔΔG_5_ = -8.1 kcal/mol (Fig. 6 purple). The actual complex is more stable than expected by -2.5 kcal/mol, due to a Gibbs coupling free energy (ΔΔG_int_) between the OXA moiety and Mg^2+^•ATP. The non-additive coupling energy between OXA and Mg^2+^•ATP does appear to be the sum of the OXA*ATP and ATP-dependent OXA*Mg^2+^ interactions (Fig. 6 blue, amber), as we were able to determine that the ΔΔG_int_ (OXA*ATP) and ATP-dependent ΔΔG_int_ (OXA*Mg^2+^) are -1.0 and -1.5 kcal/mol, respectively.

**Figure 6:**
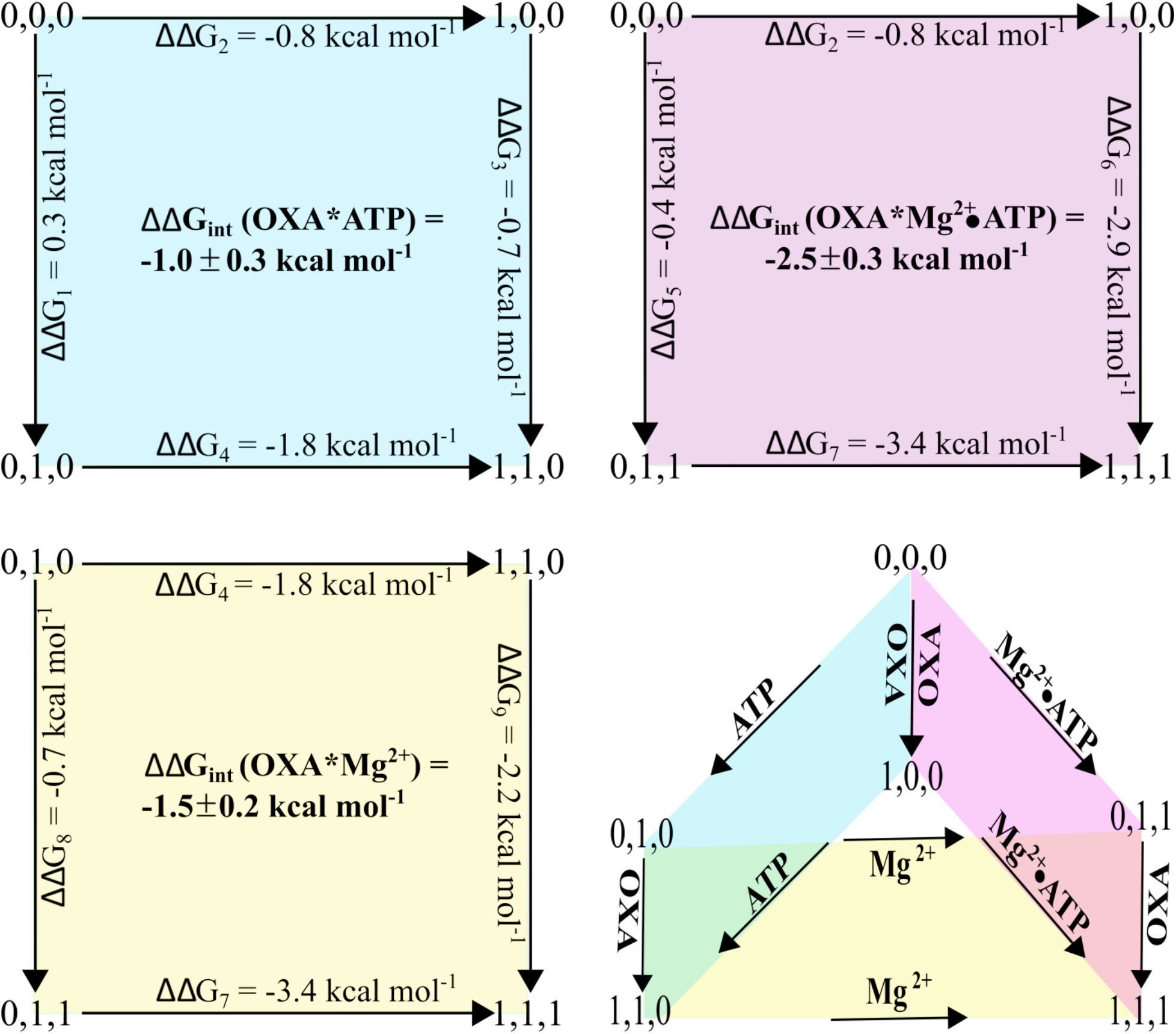
Non-additive contributions of OXA, ATP and Mg^2+^ to the thermodynamics of indolmycin inhibition. The structural differences between tryptophanamide and indolmycin account for a difference in Gibbs free energy (ΔΔG_2_) of -0.8 kcal/mol. This stabilizing effect more than quadruples to -3.4 kcal/mol (ΔΔG_7_) in the presence of Mg^2+^•ATP due to an interaction energy (ΔΔG_int_) between the methylamino-substituted oxazolinone moiety and Mg^2+^•ATP of -2.5 kcal/mol. (a,b,c) = (OXA, ATP, Mg^2+^); 0 = absent; 1 = present; tryptophanamide was the titrant when a=0; indolmycin when a=1.

The thermodynamic cycles depicted in Figure 6 validate predictions 2 and 3 described in a previous section. A consequence of our expectation that ATP is required for Mg^2+^ binding to the active site of H_mt_TrpRS (prediction 1) is that the third component (Mg^2+^) is dependent on the second component (ATP). This dependence collapses the cube resulting from testing all possible two-way interactions with three variables to a three-sided prism (Fig. 6 prism). Figure 6 (blue) proves that despite a lack of direct contacts between ATP and indolmycin in the crystal structure, a coupling interaction between these two ligands stabilizes the inhibited complex by -1.0 kcal/mol. In the absence of the metal this interaction must be mediated through the enzyme.

Finally, the addition of Mg^2+^•ATP favors indolmycin binding by ΔΔG_6_ = -2.9 kcal/mol (prediction 3) (Fig. 5b, 6 purple). ATP only accounts for 24% of this effect (ΔΔG_3_ = -0.7 kcal/mol), with the overwhelming majority of the stabilization energy coming from the Mg^2+^ ion (ΔΔG_9_ = -2.2 kcal/mol) (Fig. 6 blue, amber).

### Mutations to active site residues have greater impact on k_cat_ than K_M_ (ATP)

To confirm the functional equivalences with BsTrpRS inferred from the H_mt_TrpRS structure, assessed the impact of amino acid substitution at positions 42, 48, 145, 181, and 229. Gln^42^, Asp^181^ and Lys^229^, which form direct polar interactions with ATP (Fig. 3c, d) and are conserved at positions 9, 146, and 195, respectively, in BsTrpRS were individually mutated to alanine while His^48^, which is analogous to BsTrpRS Thr^15^, was mutated to threonine. Although we did not observe direct interactions with ATP, we chose to substitute Lys^145^ with an alanine due in part to the structural variation between monomers A and B as well as the 18.5-fold decrease in k_cat_ observed for the corresponding K111A BsTrpRS mutant (7).

These amino acid substitutions cause less than a two-fold change in ATP affinity but exhibit a 230-fold range in their impact on k_cat_ (Fig. 7, Table 4). The H48T variant retained the highest catalytic proficiency (k_cat_/K_M_) with a modest 1.4-fold reduction in k_cat_ while substitution of alanine for Lys^145^ eliminated ∼75% of the enzyme’s catalytic proficiency. By comparison, the K111A BsTrpRS variant is 94% less profficient than wild-type BsTrpRS. Alanine substitution of Gln^42^, which forms a hydrogen bond with both non-bridging alpha-phosphate oxygen atoms of ATP, preserves the weak backbone interaction (3.4 Å) but eliminates the hydrogen bond formed between the side chain amide nitrogen and the metal-coordinated P_α_ oxygen (3.0 Å). The Q42A mutant has a relative catalytic proficiency of 0.15 due primarily to a 9.4-fold reduction in k_cat_. Alanine substitution of either Asp^181^ or Lys^229^ disrupts the electrostatic environment of the active site as well as the hydrogen bonding network between the enzyme and ATP substrate. Removal of the negatively-charged Asp^181^ results in a 50-fold reduction in k_cat_. Alanine substitution of Lys^229^ has an even greater effect on catalysis. The catalytic efficiency of the K229A mutant enzyme is only 0.5% that of wild-type H_mt_TrpRS. Thus, as observed both for BsTrpRS (7) and BsTyrRS (26), active-site interactions with ATP appear to manifest mutational effects only in transition state affinity.

**Table 4:** Reduced catalytic efficiency of active site mutants is driven by diminished k_cat_. Amino acid substitutions to residues involved in ATP binding to H_mt_TrpRS have a broader effect on k_cat_ than K_M_ ATP. Alanine substitution at either position 181 or 229 abolished ∼99% of the catalytic efficiency while the K145A and Q42A variants lost 75% and 85%, respectively.

| **H<sub>mt</sub>TrpRS** | **$k_{\text{cat}}$ (s<sup>-1</sup>)** | **$K_{\text{M}}$ (M)** | **$k_{\text{cat}}/K_{\text{M}}$ (s<sup>-1</sup> M<sup>-1</sup>)** |
| --- | --- | --- | --- |
| WT | 4.61 ± 0.03 | 3.59E-04 ± 2.23E-05 | 1.29E04 ± 7.34E02 |
| H48T | 3.28 ± 0.01 | 3.95E-04 ± 3.00E-06 | 8.30E03 ± 5.08E01 |
| K145A | 1.27 ± 0.13 | 4.38E-04 ± 1.24E-04 | 3.09E03 ± 1.09E03 |
| Q42A | 0.49 ± 0.01 | 2.55E-04 ± 2.52E-06 | 1.92E03 ± 2.05E01 |
| D181A | 0.09 ± 0.01 | 5.36E-04 ± 9.09E-05 | 1.60E02 ± 1.49E01 |
| K229A | 0.02 ± 0.01 | 3.71E-04 ± 7.33E-05 | 6.35E01 ± 2.37E01 |

**Figure 7:**
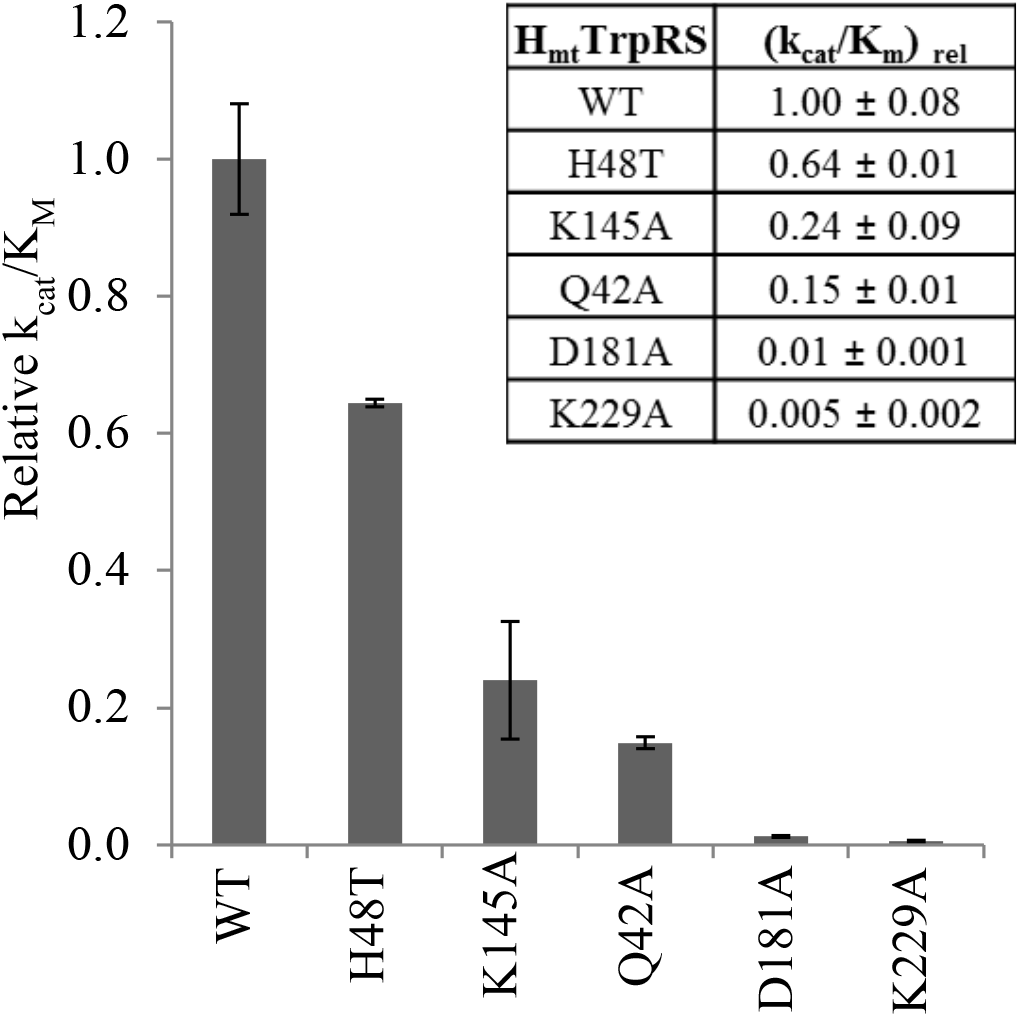
Mutations to active site residues have varying effects on catalytic efficiency. Data from steady-state Michaelis-Menten experiments were fitted by non-linear regression modeling to determine k_cat_ and K_M_ ATP for six H_mt_TrpRS variants. The catalytic efficiency of each variant was determined and normalized to the wild-type Δ1-18 enzyme. The H48T and K229A mutants were the most and least efficient variants, respectively.

## DISCUSSION

Distinctive evolutionary divergences between the eukaryotic and bacterial TrpRS enzymes (31, 32) account for a ∼10^6^-fold difference between their relative sensitivity to competitive inhibition by the natural tryptophan analog, indolmycin. The crystal structure of human mitochondrial TrpRS, together with thermodynamic and kinetic data in Figs. 2-6 and Tables 3,4 and previous mechanistic studies of BsTrpRS (6-8) and its ternary complex with ATP and indolmycin (5) enable us to describe with some confidence the basis for this extraordinary difference in relative sensitivity. Indolmycin interacts with Mg^2+^•ATP to form a unique, off-path, ground-state coordination of the catalytic metal that reduces its access to the activated configuration necessary to assist catalysis of tryptophan activation. That effect strengthens indolmycin binding by ∼1000-fold in bacterial TrpRSs.

Curiously, there is low conservation between the cytosolic, mitochondrial and bacterial TrpRS homologs at residues shown to form stabilizing interactions with tryptophan in H_c_TrpRS (Fig. 1). These sequence and structural variations affect the way the respective enzymes respond structurally following substrate binding, how they couple conformational changes to catalysis, the manner in which they retain the tryptophanyl-5’ AMP intermediate, and their ability to discriminate between tryptophan and other indole-containing compounds. This latter feature protects eukaryotic cytosolic TrpRS enzymes from indolmycin inhibition by ∼1000-fold while, as will be discussed, leaving mitochondrial and most prokaryotic TrpRS enzymes susceptible to inhibition.

### Shared inhibition mechanism of BsTrpRS and H_mt_TrpRS by indolmycin

Structural, kinetic and thermodynamic studies of Bs and H_mt_ TrpRSs identify a shared mechanism for indolmycin inhibition of prokaryotic and mitochondrial TrpRS enzymes. Specifically, a conserved aspartic acid (Bs Asp^132^; H_mt_ Asp^167^) in the specificity helix recognizes the indole nitrogen while recognition of the methylamino-nitrogen and carbonyl oxygen atoms of indolmycin by ND1 of His^77^ (Bs His^43^) and the side chain amide nitrogen of Gln^182^ (Bs Gln^147^), respectively, restrict the plane of the oxazolinone ring. Indolmycin is further stabilized by hydrogen bonding interactions via the carbonyl oxygen and oxazolinone nitrogen atoms with two metal-coordinated water molecules (Fig. 3a, b).

We have used different thermodynamic measurements—Thermofluor with BsTrpRS and ITC here—to show that the catalytic metal ion is required for the non-additive interaction energy between indolmycin and ATP (5). The results measured by the two methods, which differ importantly in the temperature at which the interaction effects are observed, are qualitatively very similar. In the presence of Mg^2+^, the oxazolinone-ATP interaction contributes ∼-2.2 kcal/mol to the indolmycin binding energy to H_mt_TrpRS. The additional contacts made with the oxazolinone moiety enable both BsTrpRS and H_mt_TrpRS to bind indolmycin with higher affinity than its tryptophan substrate while the oxazolinone•Mg^2+^•ATP interaction appears to lock both enzymes in a stable, off-path inhibited state, enhancing indolmycin affinity.

### A conserved aspartic acid recognizes indole nitrogen

Human mitochondrial TrpRS has an aspartic acid residue in its specificity helix that is conserved among prokaryotic but not eukaryotic cytosolic TrpRS enzymes. The side-chain of this residue is used by prokaryotic TrpRSs to recognize the indole nitrogen of the tryptophan substrate and contributes to binding stability via a hydrogen bond. Eukaryotic cytosolic TrpRS enzymes, which lack this aspartic acid residue, rely on a tyrosine and glutamine that are closer to the N-terminus for recognition of the indole nitrogen (31). H_c_TrpRS, which lacks a homologous aspartate in its specificity helix, undergoes induced fit rearrangement of its active site in response to tryptophan, rather than ATP, binding (9-11). This rearrangement relies on specific hydrogen bonding between the tryptophan substrate and active site residues, allowing H_c_TrpRS to better discriminate between tryptophan and other indole-containing compounds. Of the six H_c_TrpRS residues that make direct contacts with tryptophan only one residue beside the strictly conserved glutamine of the GxDQ motif is conserved in H_mt_TrpRS (Fig. 1). Tryptophan binding to BsTrpRS, promoted by a single residue-specific hydrogen bond between the indole nitrogen of tryptophan and Asp^132^ of the specificity helix supplemented by a hydrophobic pocket, does not induce structural changes of the active site (23).

### H_mt_TrpRS binds both tryptophan and indolmycin ∼40-fold more weakly than BsTrpRS

Despite differences in binding determinants and structural consequences of tryptophan binding, both BsTrpRS and H_c_TrpRS bind tryptophan with comparable binding affinities of ∼2 uM. In contrast, H_mt_TrpRS, which has a high degree of similarity between active-site residues with BsTrpRS, binds both tryptophan and indolmycin about an order of magnitude more weakly than does BsTrpRS. Examination of the tryptophan-binding pocket for the three enzymes did not reveal any obvious reason for the weaker binding to H_mt_TrpRS. Residues in this area are highly conserved between BsTrpRS and H_mt_TrpRS, as they use the same determinants for tryptophan recognition (Asp and van der Waals interactions). The only difference we found that may account for the observed differential binding affinities is the replacement of a methionine in the specificity helix for a glutamine in H_mt_TrpRS. In BsTrpRS the methionine sulfur atom is in position to interact with the aromatic ring of substrate tryptophan. H_c_TrpRS has a glutamine instead of methionine at this position but likely compensates for possible lost sulfur-aromatic interaction via numerous hydrogen bonding interactions that were not observed for either BsTrpRS or H_mt_TrpRS (33).

### Role of active-site lysines in ATP binding and catalysis

The importance of positively charged active site lysine residues in stabilizing the negatively-charged triphosphate oxygen atoms of ATP has been studied extensively in the case of BsTrpRS (7, 23, 34). Lysines 111 and 192 compete with the catalytic Mg^2+^ ion for stabilizing the negative charge of the PPi leaving group, while Lys^195^ interacts with a non-bridging alpha-phosphate oxygen as well as the oxygen atom bridging the alpha and beta phosphates.

From the fully-liganded, inhibited H_mt_TrpRS complex reported here, it is evident that ATP binds in a conformation similar to that observed in all ATP-bound BsTrpRS structures with a catalytic metal making electrostatic interactions with oxygen atoms from each phosphate group (5, 23). Lysine^226^ (BsTrpRS Lys^192^) competes with the metal for the same beta-phosphate oxygen, while Lys^229^ (BsTrpRS Lys^195^) forms electrostatic and hydrogen bonding interactions with an alpha-phosphate oxygen atom. Both interactions mimic comparable ones observed in BsTrpRS. Indolmycin binding weakens lysine-ATP interactions while strengthening ATP-metal interactions. Part of a highly mobile loop the positioning of Lys^111^ varies depending on the ligands bound to the active site. The strength of the interaction with gamma-phosphate oxygen is weaker in the presence of indolmycin than tryptophanamide (and presumably tryptophan). This residue (Lys^145^) is conserved in H_mt_TrpRS but its structure varies between the two monomers that make up the dimer in the crystallographic asymmetric unit. While neither lysine is close enough for direct interactions with ATP in the inhibited complex, our kinetic characterization of a K145A mutant (Fig. 7, Table 4) shows that this residue contributes to the stability of the catalytic transition state as was observed for Lys^111^ in BsTrpRS (7). Together, these data point to a shared role of active site lysines in the binding of ATP as well as the neutralization of negatively charged PPi leaving group following tryptophan activation between BsTrpRS and H_mt_TrpRS. However, the contribution of Lys^145^ to transition state stabilization of H_mt_TrpRS (Table 4) is not as great as that of Lys^111^ for BsTrpRS. H_c_TrpRS is missing lysines at analogous positions for both 111 and 195, the consequence of which is altered ATP configuration (bent instead of extended) and Mg^2+^•ATP coordination.

### Implications for tryptophanyl-5’ AMP formation by H_mt_TrpRS

Differences in the relative positioning of the tryptophan and ATP binding sites result in different conformers of tryptophanyl-5’ AMP (TYM) occupying the BsTrpRS and H_c_TrpRS active sites (10, 30). The TYM bound to BsTrpRS is more extended, with the indole moiety CE3 atom being 1.6 Ȧ further away from the 3’OH group than in the corresponding TYM of H_c_TrpRS. Despite these differences both H_c_TrpRS and BsTrpRS retain the enzyme-indole hydrogen bonding interactions during catalysis for continued assurance that the bound substrate is tryptophan. H_mt_TrpRS should likewise preserve the indole-Asp^167^ interaction with the TYM product.

Rigid-body fitting of the BsTrpRS TYM product into the H_mt_TrpRS active site by superposing both the indole and adenine rings maintains the Asp^167^-indole nitrogen interaction and other interactions with the AMP moiety, including those with Asn^51^ and Gln^42^ without introducing steric clashes. In contrast, rigid-body fitting of the TYM from H_c_TrpRS into the H_mt_TrpRS active site by aligning either the indole or adenine rings results in clashes with numerous active-site residues. Additionally, aligning the H_c_TrpRS TYM to the adenine moiety of ATP eliminates the Asp^167^-indole interaction. This analysis suggests that H_mt_TrpRS is likely to go from a preTS to product state via a transition mimicking that of BsTrpRS, with the TYM product adopting the more extended conformation observed in the BsTrpRS active site (23, 30).

### Mg^2+^-H_mt_TrpRS coupling

The similar ATP binding conformation and ATP-metal coordination observed for H_mt_TrpRS and BsTrpRS as well as the lack of metal-protein interactions raise again questions about metal-enzyme coupling (35). Specifically, is the catalytic metal coupled to a site remote from the H_mt_TrpRS active site as has been demonstrated for the BsTrpRS enzyme? The D1 switch identified in BsTrpRS consists of Ile^4^, Phe^26^, Tyr^33^, and Phe^37^. Re-packing of these side-chains is coupled to movement of the metal into a catalytically-competent position and D1-Mg^2+^ coupling accounts for the increased tryptophan specificity over tyrosine of BsTrpRS primarily through effects on k_cat_ (8, 36). Other work has identified repacking of the four corresponding BsTrpRS side chains with a conformational transition state limiting the rate of induced-fit (25). Structure-based sequence alignment of the inhibited BsTrpRS and H_mt_TrpRS structures identified the analogous H_mt_TrpRS residues as Val^37^, Trp^59^, Tyr^66^, and Tyr^71^ (Fig. 1), preserving all three aromatic side chains as well as the branched beta carbon of the N-terminal residue. Additional structures along the catalytic path are needed to identify side-chain rearrangements (re-packing) that occur as a consequence of substrate binding and catalysis in H_mt_TrpRS.

In this work, we confirmed our hypothesis that conserved elements, including Asp^167^ and His^77^, leave H_mt_TrpRS vulnerable to inhibition by indolmycin. The 10^-8^ M inhibition constant is an order of magnitude weaker than that measured for BsTrpRS, consistent with K_M_ tryptophan values for the two enzymes (5). The 1.82 Å structure and the associated thermodynamic studies obtained by isothermal titration calorimetry validated the essentials of our proposed mechanism by which indolmycin inhibits BsTrpRS, specifically by simultaneously weakening enzyme-ATP and strengthening metal-ATP interactions to prevent metal activation (5).

The structural data suggests that indolmycin binding should be tighter in the presence of Mg^2+^•ATP. Isothermal titration calorimetry experiments validated this claim as pre-binding Mg^2+^•ATP enhanced indolmycin binding to H_mt_TrpRS ∼100-fold, whereas ATP alone lead to a three-fold increase in indolmycin affinity. Finally, this sole structure of H_mt_TrpRS allowed us to determine that it employs conserved residues and structural elements to recognize the NE1 indole atom and to bind ATP in an extended conformation. Both of these features result in an active site arrangement mimetic of BsTrpRS but that varies substantially from that of H_c_TrpRS. Together, work presented here imply a shared mechanism for tryptophan activation between H_mt_TrpRS and BsTrpRS.

## Nota bene

This manuscript was submitted to *Journal of Biological Chemistry* by Tishan Williams (TW) in or around 2017 based on her doctoral thesis (37). Reviewers asked for minor modifications for clarity and TW never responded to those concerns. Recent interest has arisen in the work it contains because of a variety of mutations that lead to Parkinson’s Disease. For that reason, CWCjr has deposited the manuscript in BioRxiv. A revision will be submitted elsewhere in due course addressing reviewers’ concerns and with modest revisions.

## Acknowledgments

This work was supported by the National Institute of General Medical Sciences, This work was supported by National Institutes of Health Grant GM90406 (to C. W. C.) and an NIGMS diversity supplement (to Tishan Williams. We are grateful to L. Kaguni for initiating this project.

## Author contributions

Tishan Williams and CWCjr conceived the experimental plan. Tishan Williams carried out all experimental work, analyzed the data and wrote the paper in consultation with CWCjr. When asked by a third party to endorse this deposition, Tishan Williams answered “Ignore”.

### The abbreviations used are

TrpRS: tryptophanyl-tRNA synthetase
Bs: *Bacillus. stearothermophilus*
Hc: human cytoplasmic
Hmt: Human mitochondrial
LTN: tryptophanamide
IND: indolmycin
PreTS: pre-transition state
TEV: tobacco etch virus
OXA: methylamino-substituted oxazolinone ring
TYM: tryptophanyl-5’AMP
ITC: isothermal titration calorimetry.

